# Divergent evolution of head morphology between marine and freshwater sticklebacks

**DOI:** 10.1101/2023.07.14.549115

**Authors:** Antoine Fraimout, Ying Chen, Kerry Reid, Juha Merilä

## Abstract

Intraspecific phenotypic differentiation is of common place occurrence, but the degree to which it reflects phenotypic plasticity or local adaptation remains often unclear. To be considered as adaptive, the differentiation must be genetically based and exceed what could be expected by neutral processes only. Using laboratory reared full-sib family data from replicate nine-spined stickleback (*Pungitius pungitius*) populations, we show that freshwater and marine fish display genetically based adaptive differentiation in head size and shape. Utilising identity-by-descent relationships among full-sibs as estimated with the aid of molecular markers, we further show that the studied traits are also highly heritable in all populations indicating and that they can respond to future episodes of natural selection. The head shape and size of pond fish suggests that observed adaptive differentiation has been driven by selection favoring limnetic feeding strategy among the pond fish. Analyses of gill-raker morphology were less conclusive: genetic differentiation was found in gill-raker length (pond > marine) and number, but the degree of divergence in these traits did not exceed neutral expectations. Yet, the direction of divergence in gill raker traits are suggestive of the limnetic feeding mode of pond fish, aligning with the inference from the head morphology analyses.

## Introduction

Phenotypic differentiation associated with different habitat characteristics is of commonplace occurrence in both animal (Endler 1978; Foster 1999) and plant (Linhart and Grant 1996; Bossdorf et al. 2005) kingdoms. Likewise, phenotypic responses to changing environmental conditions have been frequently reported (Walther et al. 2002; Parmesan 2006; Sheridan & Bickford 2011), but many studies struggle to disentangle phenotypic plasticity and genetically based differentiation as the underlying cause for observed differentiation (Gienapp et al. 2008; Merilä 2012; Merilä & Hendry 2014; Stamp & Hadfield 2020). Moreover, even if evidence for genetically based differentiation is found, additional evidence is required to prove that the observed differentiation is adaptive.

Common garden studies combined with neutrality tests (e.g., Lande 1976; Rogers 1986; Spitze 1993; Merilä & Crnokrak 2001) provide a fairly straightforward way to differentiate adaptive *vs.* non-adaptive causes of phenotypic differentiation (Whitlock 2008; Leinonen et al. 2013; Savolainen et al. 2013). In particular, the neutrality tests implemented in the program ‘driftsel’ (Karhunen et al. 2013, 2014) provide a statistically powerful approach able to identify footprints of natural selection even in a fairly small sample of populations (Ovaskainen et al. 2011). Nevertheless, the driftsel-approach has not become widely used in studies of population differentiation, possibly because the the field has moved to use genomic data to identify footprints of natural selection (e.g. Storz 2005; Nosil et al. 2009; Narum & Hess 2011), and because common garden studies are logistically demanding. While genomic studies of natural selection appear to provide less laborious paths towards identifying adaptive differentiation than common garden experiments, they are still grappling with the problem of false positive tests (Mallick et al. 2009; Bierne et al. 2013; Hoban et al. 2016), as well as with the difficulty of linking the loci under selection to their phenotypic targets especially in the case of polygenic traits (McKay & Latta 2002; De Kovel 2006; Le Corre & Kremer 2012). In addition, genome scan approaches become very inefficient in picking up signals of selection in systems having high background levels of differentiation, such as in highly subdivided populations experiencing strong drift (Hoban et al. 2016).

Variation in fish head shape and jaw structures have been extensively studied as they are important trophic traits influencing fitness (e.g., Roy et al. 2010; McGee et al. 2013) and even believed to have spearheaded adaptive radiations (Brouwers 2011; Sallan & Friedman 2012). While population differences in them have been sometimes shown to be genetically based (Kimmel et al. 2005; Albert et al. 2008; McGee & Wainwright 2013), there is also evidence that there is a strong plastic component to variation in head shape and feeding structures (Troy et al. 1994; Dingemanse et al. 2009). Hence, while there are many reports of head and jaw shape differentiation among fish populations (e.g., Walker and Bell 2000; Tobler et al. 2011 Østbye et al. 2016), few studies have firmly established an adaptive (contra plastic) basis for this differentiation.

Gill-raker morphology constitutes another set of traits that have been extensively studied in the context of fish foraging ecology (e.g., Berner et al. 2008; Wund et al. 2012; Hosoki et al. 2019). Fish species have adapted to various habitats by evolving unique gill-raker characteristics that correspond to their specific feeding strategies. As for the case of three-spined sticklebacks (*Gasterosteus aculeatus*), marine populations typically have more, closely spaced and longer gill rakers than freshwater populations (Gross and Anderson. 1984; Leaver et al. 2012; Glazer et al. 2014; Magalhaes et al. 2021), reflecting their limnetic feeding strategy in marine ecosystems and the benthic feeding strategy in freshwater ecosystems (Ravinet et al. 2014). However, formal tests of the adaptive basis of this differentiation are few (but see: Raeymaekers et al. 2007; Seymour et al. 2019).

The aim of this study was to test (1) whether pond and marine populations of nine-spined sticklebacks (*Pungitius pungitius*) differ in head and gill raker morphology and (2) whether this differentiation exceeds neutral expectations meaning that observed divergence has been driven by natural selection. To do this, we conducted a common garden experiment and raised fish from four pond and four marine populations to control for environmental effects on phenotypes, and subjected the data to driftsel analyses to identify footprints of selection. The results revealed genetically based phenotypic differentiation in head and gill-raker traits between marine and pond sticklebacks, and that the differentiation in head traits has been driven by divergent natural selection.

## Methods

### Study populations and sampling

The parental fish used in this study were collected May-June 2018 from four pond (Rytilampi 66.38482°N, 29.31561°E; Pyöreälampi 66.26226°N, 29.42916°E; Kirkasvetinenlampi 66.43673°N, 29.13568°E; Bynastjärnen 64.45416°N, 19.44075°E) and four marine (Tvärminne 59.83333°N, 23.24900°E; Pori 61.59111°N, 21.47295°E; Raahe 64.68818°N, 24.46189°E; Bölesviken 63.66110°N, 20.20940°E) populations from Fennoscandia using minnow traps or beach seine (Fig. 1a). The captured adults were transported to aquaculture facilities of University of Helsinki, and maintained in 1 m^3^ flow-through freshwater aquaria (one per population) until used in artificial fertilisations.

**Fig. 1.**
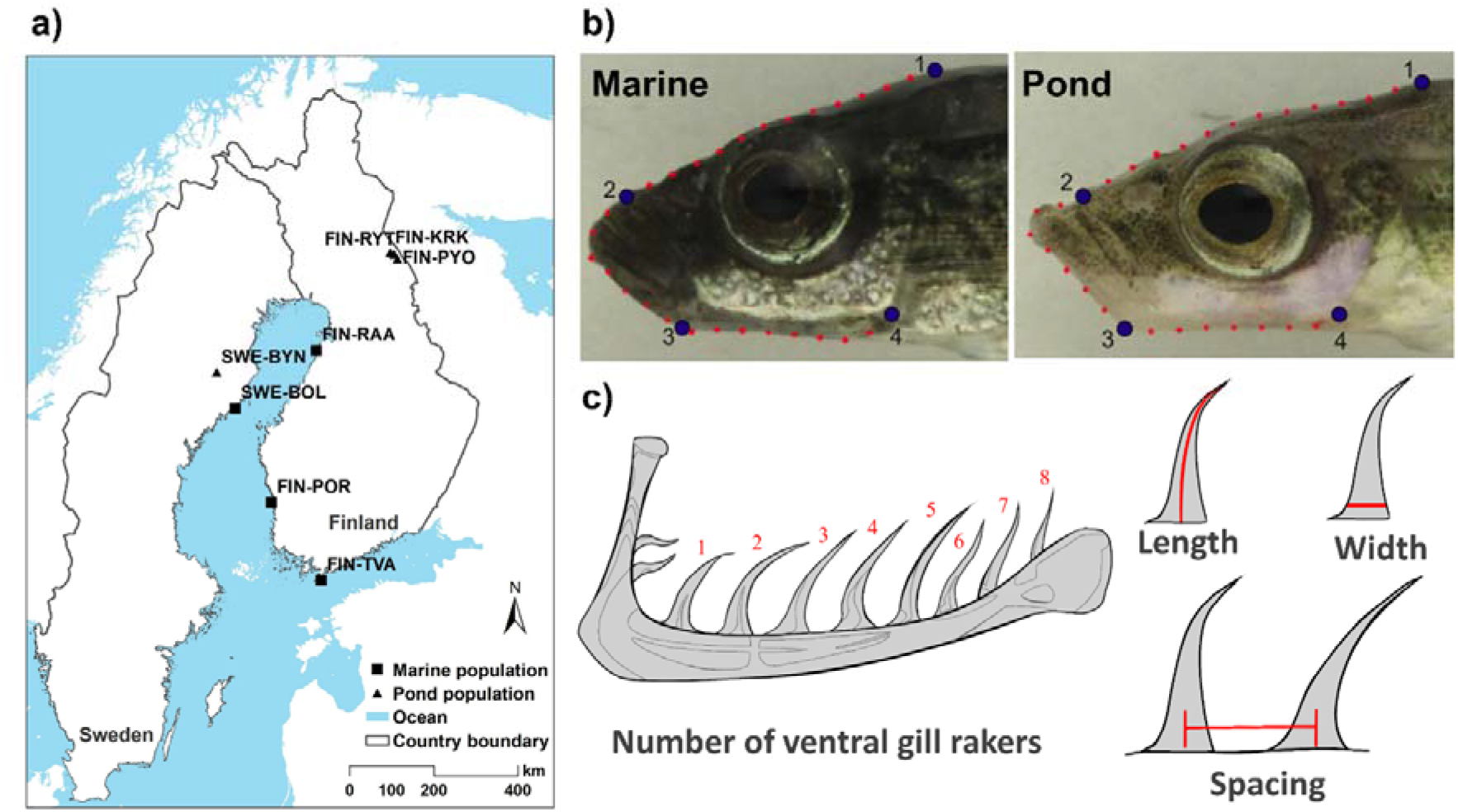
a) Map of the study area displaying the eight sampling locations of nine-spined stickleback populations, which include four marine (Tvärminne, TVA; Pori, POR; Raahe, RAA; Bölesviken, BOL) and four pond populations (Rytilampi, RYT; Pyöreälampi, PYO; Kirkasvetinenlampi, KRK; Bynastjärnen, BYN) in Finland and Sweden. b) Landmark positions. Positions of the four fixed landmarks (blue filled circles) and semilandmarks (red filled circles) used to quantify head shape variation in marine (left panel) and pond (right panel) populations. Numbering of fixed landmarks are as in *Methods*. c) Diagram illustrating the four gill raker trait measurements: number, length, width, and spacing between rakers.

### Common garden experiments and genotype data

Artificial fertilizations among randomly selected adult fish in breeding condition were conducted between May 24^th^ and July 4^th^ 2018. Standard *in vitro* fertilization techniques and egg rearing methods were applied following Arnott and Barber (2000) and as described in detail in Fraimout *et al*. (2022). In brief, eggs from gravid females were obtained by gently squeezing their abdomens over a petri dish. Sperm was retrieved from males after they were over-anesthetized with tricaine methanesulfonate (MS-222) by dissecting the testes and subsequently mincing them in the same petri dish containing the eggs. Eggs and sperm were mixed to ensure fertilization and kept in water until hatching. To avoid fungal infections, water in the petri dishes was changed twice daily and clutches were inspected for signs of fungal infections or death - dead and infected eggs were removed. At hatching each clutch was split in two replicate 11 x 10 x 10 cm plastic containers filled with filtered freshwater in which they were maintained for a four week period during which yolk resorption took place. The larvae were fed *ad libitum* with live brine shrimp (*Artemia sp*. nauplii). After this, all replicated families were transferred to Allentown Zebrafish Rack Systems (Aquaneering Inc., San Diego, USA). These racks had a closed water circulation system with physical, chemical, biological and UV filters. The fish in racks were fed for the first four weeks with a mixture of *Artemi*a nauplii and chopped chironomid larvae, and after this, with chopped chironomid larvae. Part of the water in the rack systems was changed every fortnight to maintain good water quality. Fish were reared in family groups (maximum five individuals per aquarium) in racks for a period of ca. 1 year (mean age: 316.4 days) after which the fish were euthanized with MS-222. Temperature and light conditions were kept constant throughout the experiment (12:12 LD; 15°C) except during an overwintering period of 3 months (November – January) when water temperature was lowered to 10°C and light cycle to 5:19 LD.

We used data on molecular variation among populations to estimate a neutral baseline for the tests of selection (see below). Specifically, we obtained genome-wide SNP data for all individuals using a restriction site-associated DNA (RAD) sequencing technique (2b-RAD; Wang et al., 2012). Genomic DNA was extracted from fin clips using a standard salting out protocol (LoperaBarrero et al. 2008) and 2b-RAD libraries were built following Momigliano et al (2018) using a slightly modified version of the protocol available online at https://github.com/z0on/2bRAD_GATK/blob/master/2bRAD_protocol_may15_2017_nnrw.pdf). Briefly, high molecular weight DNA was digested using the *BcgI* enzyme (New England Biolabs), adaptors were ligated and the fragments amplified via PCR following the protocol provided in Momigliano *et al*. (2018). Libraries were pooled and the target fragments were isolated using a BluePippin size selector (Sage Science). Libraries were sequenced using Illumina technology (HiSeq 4000) at the Beijing Genomic Institute (BGI; Hong-Kong). Raw reads were demultiplexed and mapped to the *P. pungitius* genome (v.6; Varadharajan *et al*. 2019) using bowtie2 (Langmead and Salzberg, 2012). We used samtools (v.1.10; Li *et al*. 2009) to convert SAM files to BAM files.

We also used SNP-data to identify sex of all F_1_ offspring, as they were measured before reaching sexual maturity. From the genotype data file, we used the *snpRelate* R package (Zheng *et al*. 2012) to perform a Principal Component Analysis (PCA) based on all markers located on the sex chromosome (Linkage group 12; Natri *et al*. 2019) and assigned sex to the F_1_ individuals according to their clustering with the parental individuals of known sex. Finally, we pruned the dataset from markers in high linkage disequilibrium (LD) using the *snpgdsLDpruning* function of the *snpRelate* package and retained only markers with LD < 0.8 and a minimum allele frequency (MAF) > 0.05 and removed all sex-linked SNPs. This resulted in a total of 2,660 informative SNPs.

### Analyses of head size and shape

We quantified variation in head size and shape among the study populations following a landmark-based geometric morphometrics approach (Bookstein 1991). A total of 32 landmarks (see Fig. 1b for details) and semi-landmarks were digitized from the left side of individuals from digital images using the tpsDig2.1 software (Rohlf 2006). Specifically, 4 fixed landmarks were placed as in Yang *et al*. (2016) at: the posterior extent of the supraoccipital (landmark 1; Fig. 1b), the anterior insertion of the premaxilla (landmark 2; Fig. 1b), the anterior-ventral extent of the preopercular (landmark 3; Fig. 1b) and the ventral extent of the preopercular bone (landmark 4; Fig. 1b). Semilandmarks were placed by resampling a total of 30 points along an outline starting from landmark 1 and ending on landmark 4 describing head shape using the ‘draw curves’ mode in tpsDig. To avoid redundancy the first and last semi landmarks (overlapping with fixed landmarks 1 and 4) were removed prior to analyses. All subsequent morphometric analyses from the obtained landmark configuration were performed in R using the *geomorph* package (v.4.0.1; Adams *et al*. 2013). First, we used the plotOutliers function in geomorph to identify outlier individuals based on their Procrustes distance from the mean head shape of the samples. We removed 16 individuals outside the upper quartile range and retained a total of 425 individuals for subsequent head shape analyses. Landmarks were superimposed following the Procrustes generalized least square superimposition (Dryden & Maria 1992) and semi-landmarks were slid to the mean shape configuration and their position optimized using Procrustes distance by setting the *ProcD* option of the *gpagen* function to “TRUE” in geomorph. Head centroid size obtained from the Procrustes alignment was used as measurement for head size in all subsequent analyses. For head shape, we performed a PCA on the procrustes coordinates using the *gm.prcomp* function. The non-null principal components (PC) describing shape variation were used as shape data in the subsequent analyses and standardized for allometric variation (Rolshausen *et al*. 2015). Specifically, we followed the allometric approach of Lleonart *et al*. (2000) where the standardized trait measurement PC_S_ is defined such that:

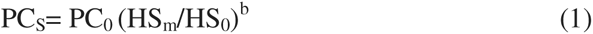

where PC_0_ is the original (i.e., unstandardized) trait measurement, HS_M_ is the mean head centroid size per population; HS_0_ is the head centroid size of each individual and b is the population-specific coefficient of within-slope regression of head centroid size on the focal PC.

We tested for population and sex differences for head size and shape using procrustes analyses of variance as implemented in the *geomorph* package. Differences in head centroid sizes were tested using the *procD.lm* function and fitting a model including body length, sex, habitat and population as explanatory variables. Similarly, we tested for differences in head shape by fitting a model including log-transformed head centroid size, sex, habitat and population as explanatory variables. We further tested for differentiation in head shape data using multivariate analysis of variance (MANOVA) based on the standardized PC described above.

### Analyses of gill raker number and size

Gill raker traits were measured from a total of 126 individuals from 64 families across eight populations — four pond (n=51) and four marine (n=75) populations. These samples included 62 wild-caught individuals and 64 of their laboratory-raised F_1_ generation offspring, comprising 32 females, 92 males, and two individuals of unknown sex.

Four gill raker traits were measured: number of gill rakers, length, width, and spacing between rakers. The number of gill rakers were counted on the left side of the first gill arch unless the left gill rakers were damaged, in which case the right side counts were used instead. As the dorsal arch is prone to damage during dissection, only gill rakers on the ventral arch were counted. We measured the length and width of the longest gill raker along the arch, and defined spacing as the center-to-center spacing between the base of a pair of adjacent rakers, specifically between the longest raker and the adjacent one on the upper side (Fig. 1c).

To ensure high repeatability of the measurements, we followed the laboratory protocol of Nicholas and Craig (2016). In summary, we first dissected the first branchial gill arch, digested it in a 10% Potassium Hydroxide (KOH) solution for two hours, and stained it with 0.008% Alizarin Red S in 1% KOH in water for at least two hours, as the staining time varies depending on the gill arch size. We then rinsed the raker to remove excess stain, placed the specimen in 89% glycerol, and flat-mounted the skeleton on bridged cover slips. Measurements were obtained using digital images taken with a fluorescent stereomicroscope equipped with an Olympus SZX16 or a Nikon FI3 camera. Length-related measurements were accurate to the nearest 0.01 mm using ImageJ software v. 1.51 (Schneider et al. 2012). To validate the measurement data, we independently measured the same set of traits for 20% of randomly selected samples and calculated the repeatability (Nakagawa and Schielzeth, 2010). The repeatability of these traits from two independent measurements ranged from 0.855 to 0.967 (p < 0.001), indicating a high level of precision in the measured traits.

As body size (standard length) varied between habitats and sexes, we size-standardized all linear gill raker traits prior to statistical analysis. This was done by constructing linear mixed effect models, incorporating habitat and sex as fixed factors and population and pedigree information as random factors, to determine if marine and pond populations differed in mean values of gill raker traits and whether these traits displayed sexual dimorphism. This analysis was performed using the ’mmer’ function in the ’sommer’ R package (Giovanny, 2016) using R version 4.2.2 (R Core Team, 2022).

### Tests of natural selection

To test whether observed population differentiation in morphology exceeded neutral expectations, the approach of Ovaskainen et al. (2011) as implemented in the R packages driftsel and rafm (Karhunen *et al*. 2013) was adopted. We followed the workflow from Karhunen *et al*. (2013) and first estimated the matrix of population-level coancestry from genomic data. Due to the computational burden of the MCMC-based algorithm, we sampled 2000 random markers from the pruned SNP dataset (Li *et al*. 2019) which were used as input for the *do.all* function of the rafm package. We ran the model for 15,000 MCMC iterations with a burn-in period of 5000 and sampling every 10th iteration. The resulting coancestry matrix was then used as a neutral baseline for the driftsel analysis. We ran the *MH* function of the driftsel package using the posterior samples of the coancestry matrix as prior information and used either head centroid size or the three first size-standardized-PC most representative of habitat shape variation as input traits. Sex of the individuals was used as covariate and body length was added as covariate for the model using head centroid size only. We subsequently used the *S.test* and *H.test* functions to calculate the *S* and *H* statistics, respectively, as indicators of signal of selection. The rationale behind each statistic is similar to that of classical Q_ST_-F_ST_ comparisons with values of *S* or *H* > 0.95 indicating signal of divergent selection, values equal to 0.5 indicates a scenario compatible with neutral evolution and values < 0.05 imply stabilizing selection. Contrary to the *S.test*, the *H.test* allows to incorporate habitat information into the driftsel framework to test whether habitat similarity correlates with phenotypic similarity among populations (Karhunen *et al*. 2014).

To verify the robustness of the selection test results, we also applied the method of Martin *et al*. (2008) to our genetic and phenotypic data. This method is a multivariate approach to the Q_ST_-F_ST_ comparison and tests for proportionality between the genetic (co)variance matrix (**G**) and the among-population divergence (**D**). Under a scenario of purely neutral evolution, the proportionality between **G** and **D** is defined as:

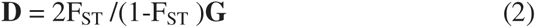

where F_ST_ is estimated from putatively neutral molecular markers (Lande 1979, Lofsvold 1988, Marroig and Cheverud, 2004, McGuiguan et al 2005, Martin et al. 2008, Berner et al 2010). We applied the framework of Martin *et al*. (2008) to calculate the coefficient of proportionality ρ_ST_ between **D** and **G**, and compared it to the value of 2F_ST_ /(1-F_ST_) obtained from the pruned SNP dataset. The rationale behind this test is that values of ρ_ST_ > 2F_ST_ /(1-F_ST_) would indicate a signal divergent selection. We used the *pairwise.prop* R function of the *neutrality* package from Martin et al. (2008) to calculate ρ_ST_ and the *hierfstat* R package (v.0.5.11; Goudet & Jombart 2022) to get bootstrapped estimates of F_ST_ from neutral loci. Statistical difference between ρ and 2F_ST_ /(1-F_ST_) was assessed based on the overlapping of confidence intervals around the two estimates.

### Heritability of head morphology

We estimated the heritability of head shape variation using animal models under a Bayesian framework using the MCMCglmm R package (Hadfield 2010). We partitioned the phenotypic variance V_P_ of head shape into its additive V_A_ genetic component and estimated heritability as h²=V_A_/V_P_. For each population, we fitted a model including the first PC (PC1) describing habitat variation in head as a response variable and added head centroid size as covariate and sex of the individuals as fixed effect. We used the same approach to estimate *h²* of head centroid size using sex of the individuals as a fixed effect. For both models, to account for actual relatedness among individuals in each population, we appended the Genomic Relationship Matrix (GRM) estimated from SNP data as a random effect in the model. The GRM was estimated from the pruned SNP data using the *G.matrix* function of the *snpReady* R package (Granatto *et al*. 2018). Each model was run with 303000 MCMC iteration with a burn-in period of 3000 and sampling every 100^th^ iteration. Model checking was performed by visually inspecting the trace plots of the MCMC chains and by inspecting the effect sizes of variance components using the *effecSize* function of the *coda* R package (Plummer *et al*. 2006). All *h²* values are reported as the median of the posterior samples along with their 95% Highest Posterior Density (HPD) intervals.

## Results

### Population differentiation in head morphology

We found significant effects of sex, population of origin and habitat on head centroid size (Table 1). Freshwater fish had larger heads than marine fish (Fig. 2a) and head centroid size varied to a greater extent among pond than among marine populations (Fig. S1). Comparison between sexes revealed sexual dimorphism in head centroid size, males having bigger relative head centroid size than females (Fig. 2a). We also found differences in head shape among habitats with the first PC of shape variation discriminating between pond and marine individuals (Fig. 2b). This difference was reflected by a significant habitat effect in the procrustes Anova on head shape (*p* <0.001, Table 2).

**Fig. 2.**
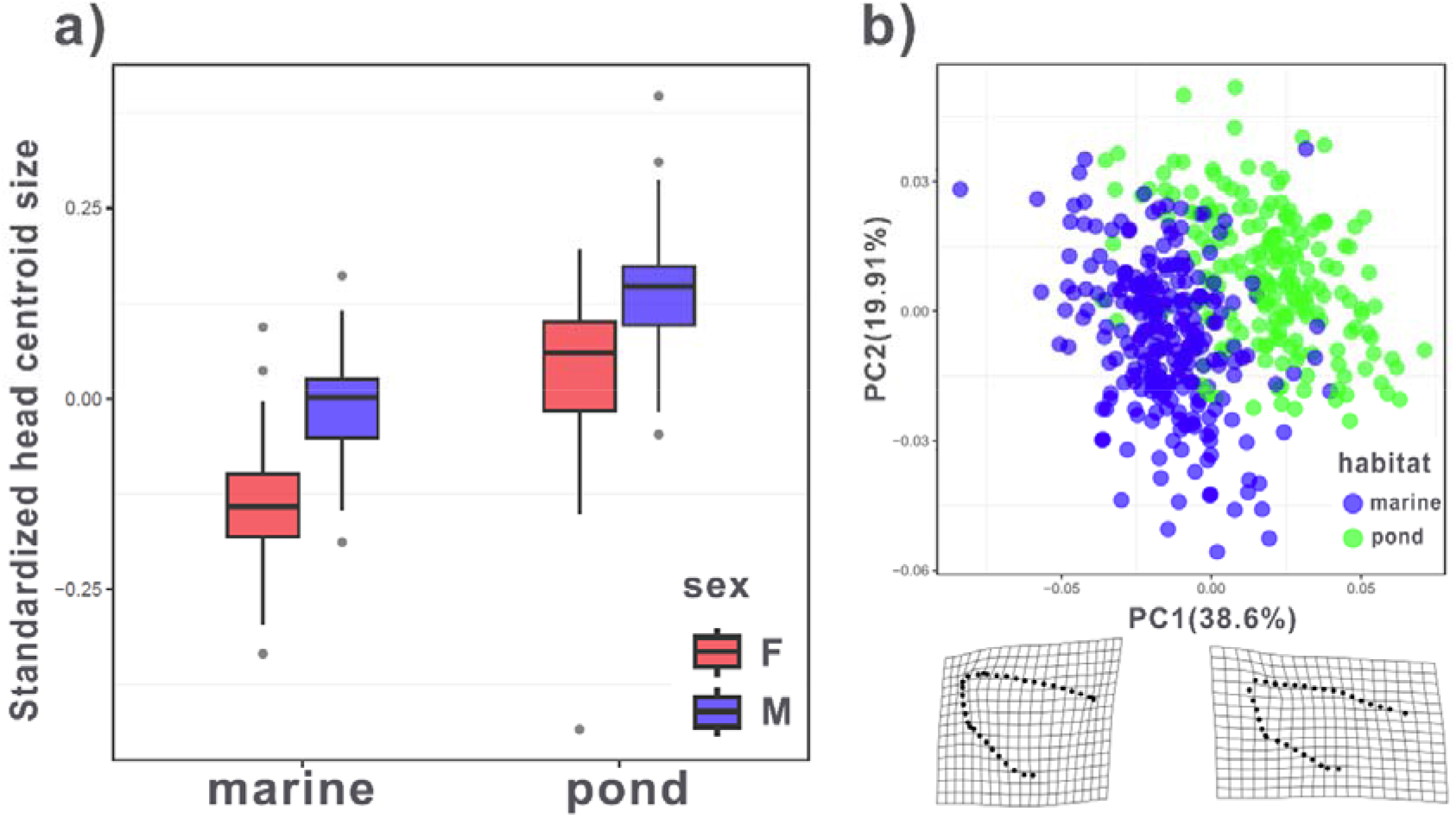
Head centroid size and shape differentiation. a) The residuals from a regression of head centroid size on body length were used to describe standardized head size and are plotted for each habitat (marine or pond) and the two sexes separately (males = M, females = F). b) Principal component (PC) analysis of head shape variation. Individuals from marine (blue filled circles) and pond (green filled circles) habitats are plotted along the two first PC axes. Deformation grids depict the shape change associated with PC1 of head shape variation.

**Table 1.** Results from the Procrustes ANOVA on head centroid size. The analysis of variance (ANOVA) table is shown for the model testing for differences in head centroid size. Effect: the fixed effect included in the models. Df: degrees of freedom. SS: sums of squares; MS: mean squares; Rsq: correlation coefficient (R²); F: values from the F-distribution; Z: effect sizes. *p*: *p-value* with bold font indicating statistical significance (*p* < 0.05).

| Effect | Df | SS | MS | Rsq | *F* | *Z* | *p* |
| --- | --- | --- | --- | --- | --- | --- | --- |
| Body length | 1 | 14.735 | 14.735 | 0.716 | 3227.578 | 12.631 | **0.001** |
| Habitat | 1 | 2.966 | 2.966 | 0.144 | 649.773 | 9.425 | **0.001** |
| Population | 6 | 0.157 | 0.026 | 0.007 | 5.737 | 3.975 | **0.001** |
| Sex | 1 | 1.424 | 1.424 | 0.055 | 250.226 | 7.018 | **0.001** |
| Residuals | 348 | 1.589 | 0.005 | 0.077 |  |  |  |

**Table 2.** Results from the Procrustes ANOVA on head shape. The analysis of variance (ANOVA) table is shown for the model testing for differences in head shape. Effect: the fixed effect included in the models with semi-columns indicating interaction. Df: degrees of freedom. SS: sums of squares; MS: mean squares; Rsq: correlation coefficient (R²); *F*: values from the F-distribution; *Z*: effect sizes. *p*: *p-value* with bold font indicating statistical significance (*p* < 0.05).

| Effect | Df | SS | MS | Rsq | F | Z | $p$ |
| --- | --- | --- | --- | --- | --- | --- | --- |
| Log(Csize) | 1 | 0.072 | 0.072 | 0.123 | 69.743 | 7.479 | **0.001** |
| Habitat | 1 | 0.085 | 0.085 | 0.144 | 81.928 | 7.176 | **0.001** |
| Population | 6 | 0.059 | 0.009 | 0.100 | 9.156 | 7.307 | **0.001** |
| Sex | 1 | 0.012 | 0.012 | 0.019 | 11.302 | 4.508 | **0.001** |
| Log(Csize):Habitat | 1 | 0.001 | 0.001 | 0.002 | 1.284 | 0.724 | 0.241 |
| Residuals | 347 | 0.359 | 0.001 | 0.610 |  |  |  |

### Population differentiation in gill raker traits

Individuals from pond populations were larger than those from marine populations (X_pond_ = 53.39 ± 2.39 [S.E.] mm, X_marine_ = 42.78 ± 1.12 mm; *F*_1,124_ = 27.75, *p* < 0.001), and females were bigger than males (X_females_ = 57.16 ± 2.52 mm; X_males_ = 43.20 ± 1.23 mm; *F*_1,123_ = 51.79, *p* < 0.001). Marine and pond populations differed significantly in both the number (X_pond_ = 9.94 ± 0.13; X_marine_ = 8.59 ± 0.08; *t*_3.02_ = 5.572, *p* = 0.01) and length (X_pond_ = 1.45 ± 0.04 mm; X_marine_ = 1.09 ± 0.02 mm; *t*_5.91_ = 2.630, *p* = 0.039) of gill rakers, as did the sexes (X_females_ = 9.11 ± 0.20; X_males_ = 9.14 ± 0.10; *t*_122.79_ = 2.794, *p* = 0.006) for number and (X_females_ = 1.31 ± 0.06 mm; X_males_ = 1.21 ± 0.03 mm; *t*_119.68_ = 4.549, *p* < 0.001) for length. Sticklebacks from pond populations had more and longer rakers (Fig. 3), and males possessed shorter rakers for their size relative to females. However, no significant differences were observed in the width or spacing of rakers across habitats or sexes (p > 0.05 in all tests).

**Fig. 3.**
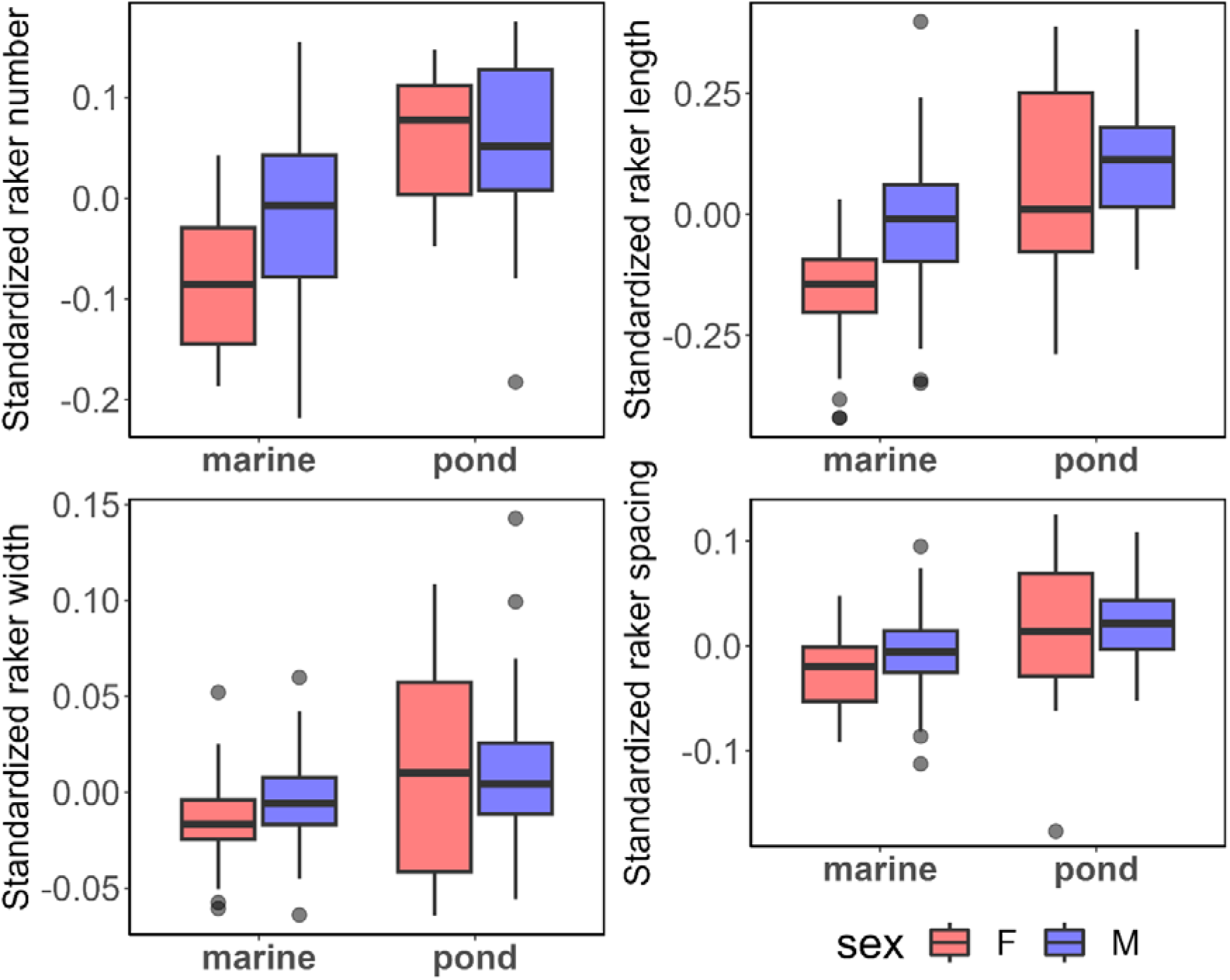
Differences in mean size standardized values of four raker traits in marine and pond environments, as well as in two sexes (females = F, males = M).

### Tests of selection

The three first PCs from the PCA of size-standardized-PC of shape variation were used as input for driftsel which revealed a signal of divergent selection (*S* = 0.953 and *H* = 0.993; Fig. 4). All populations from both habitats were indicated to have diverged more than expected by chance from their common ancestor (Fig. 4). The direction of divergence differed consistently for marine and pond populations, with the exception one pond population (KRK; Fig. 4b). Individuals from this population had an intermediate head morphology compared to other populations and were not significantly different from the reconstructed ancestral population (Fig. 4). We found similar results when analyzing head centroid size differentiation with *S* = 0.978 and *H* = 1.

**Fig. 4.**
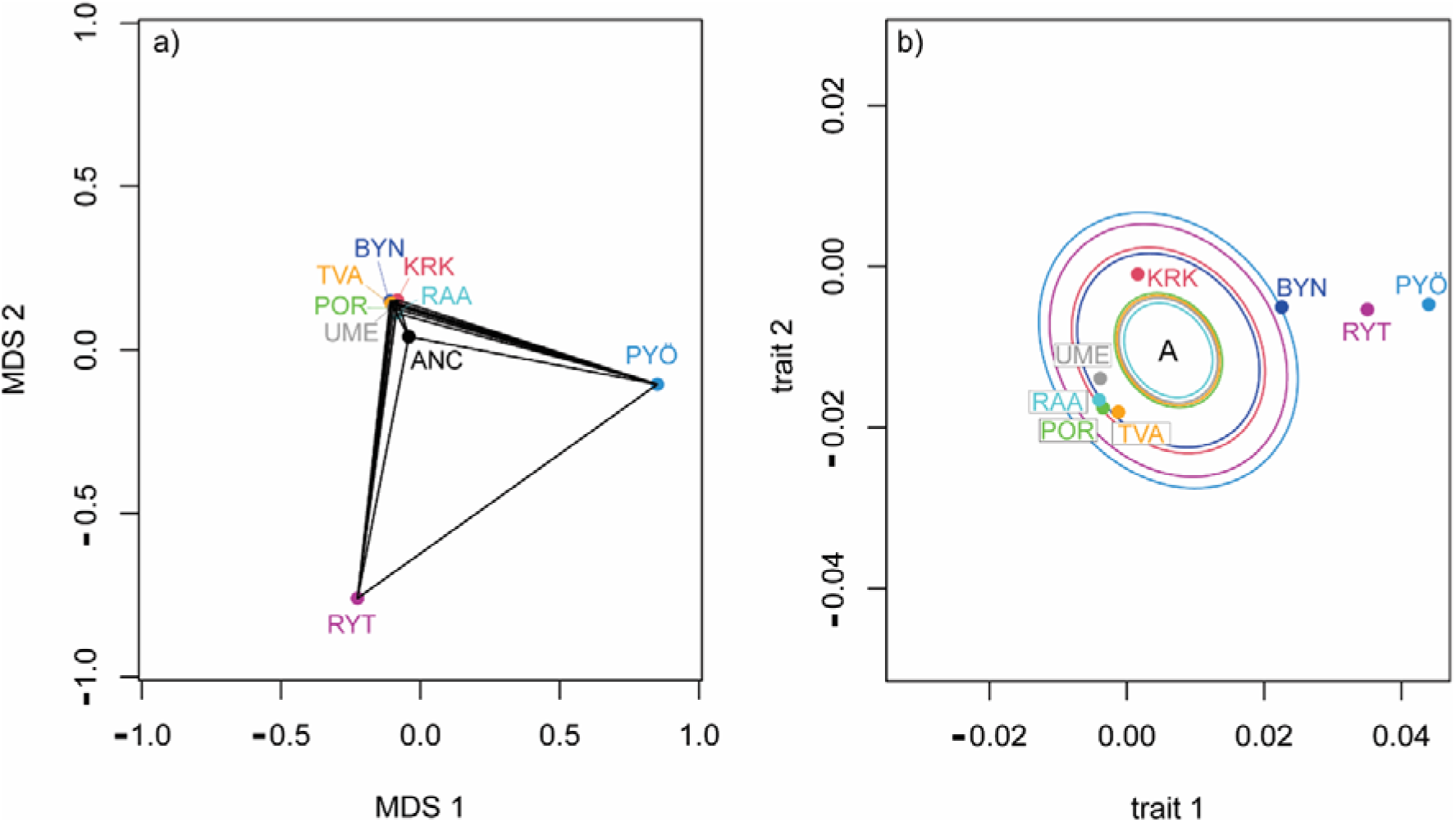
Results of the driftsel analysis. a) Multidimensional scaling (MDS) patterns of neutral genetic differentiation estimated from SNP data obtained from the *viz.theta* function in program driftsel. b) Population means in the trait space from the *viz.trait* function. Ellipses represent the expected median distance from the inferred ancestral population (ANC) under neutral evolution.

The multivariate Q_ST_-F_ST_ approach indicated that the proportionality coefficient ρ was higher than 2F_ST_ /(1-F_ST_) obtained from neutral markers (ρ_ST_ = 1.8 [1.1; 5.3]; 2F_ST_ /(1-F_ST_) = 0.828 [0.766; 0.890]), thus indicating a signal of divergent selection for head shape. Neither X² or Bartlett-corrected X² test rejected the null hypothesis of proportionality between **D** and **G** (*p* = 0.07; *p*=0.159, respectively) thus indicating that both matrices are proportional.

We did not find evidence for diversifying natural selection for any gill raker trait (*S* = 0.58, *H* = 0.86 for the univariate model for raker number; *S* = 0.69, *H* = 0.89 for length; *S* = 0.49, *H* = 0.62 for width; and *S* = 0.54, *H* = 0.74 for gap), suggesting that pond and marine populations have not evolved significantly further away from the ancestral mean than would be expected under neutrality. The same conclusion was reached combining all four gill raker traits in a multivariate analysis (*S* = 0.55; *H* = 0.91). This was also confirmed by the proportionality coefficient (ρ_ST_ = 0.33 [0.21; 0.80]) being not significantly higher than 2F_ST_ /(1-F_ST_) of 0.828 [0.766; 0.890]).

### Heritability of head size and shape

We found moderate to high *h²* values for head shape and size in all populations (Table 3). Estimates for different populations were not significantly different as indicated by the overlapping 95% HPD intervals (Table 3).

**Table 3.** Estimates of heritability for head centroid size and head shape. For each population, the posterior median of *h²* is shown for head centroid size (*h²*_size_) and the PC1 of head shape variation *(h²*_shape_) along with their 95% highest posterior density intervals.

| Population | $h^2_{\text{size}}$ | $h^2_{\text{shape}}$ |
| --- | --- | --- |
| SWE-BYN | 0.466 [0.223; 0.766] | 0.566 [0.393; 0.718] |
| FIN-KRK | 0.509 [0.168; 0.826] | 0.519 [0.286; 0.778] |
| FIN-PYO | 0.449 [0.222; 0.723] | 0.543 [0.363; 0.721] |
| FIN-RYT | 0.389 [0.125; 0.701] | 0.674 [0.538; 0.785] |
| SWE-BOL | 0.464 [0.221; 0.732] | 0.529 [0.387; 0.655] |
| FIN-RAA | 0.424 [0.201; 0.682] | 0.630 [0.489; 0.741] |
| FIN-TVA | 0.504 [0.250; 0.270] | 0.562 [0.439; 0.700] |
| FIN-POR | 0.499 [0.221; 0.798] | 0.559 [0.415; 0.710] |

## Discussion

The results revealed consistent differences in head and gill-raker morphology between marine and pond populations of nine-spined sticklebacks reared in common garden conditions. This together with the results of the selection tests provide firm evidence that the genetically based differentiation in head size and shape (but not in gill-raker morphology) has been driven by divergent natural selection favouring different head morphology in marine and pond habitats. Since the studied pond populations have evolved independently in total isolation since their colonisation after the last glaciation (Herczeg et al. 2009; Shikano et al. 2010), the results also provide strong evidence for parallel phenotypic evolution in head morphology: random processes are not expected to result in evolution of similar morphologies in different localities (Schluter 2000). However, even if the heritabilities of head traits were similar in different populations, the question whether the genetic underpinnings of the similar head morphology in different pond populations are underlied by the same or different genetic loci remains to be investigated. Earlier studies of other traits in these populations suggest that the similar phenotypic outcomes have been the result of recruitment of different genetic loci to underlie similar phenotypic adaptations (Kemppainen et al. 2021; Fraimout et al. 2022; Yi et al. 2023). In the following sections, we discuss how the results advance our understanding of local adaptation in general, and that of sticklebacks in particular.

The driftsel approach circumvents many shortcomings associated with traditional Q_ST_-F_ST_ approaches aiming to test whether trait differentiation among populations exceeds that to be expected due to stochastic processes alone (Karhunen et al. 2013). Here, it provided evidence for adaptive nature of head size and shape differentiation among pond and marine stickleback populations. As such, the results align with earlier evidence for adaptive differentiation among the studied pond and marine populations (e.g. Herczeg et al. 2009; Ab Ghani et al. 2013; Karhunen et al. 2014; Fraimout et al. 2022). Particularly noteworthy is that despite the high degree of background differentiation (mean F_ST_ = 0.286 across all populations) in our data, driftsel was able to pick up a signal of divergent natural selection. The approach of Martin *et al*. (2008) also suggested that differentiation in head shape was greater than the neutral expectation but that the magnitude of divergence was moderate. Nevertheless, it is conceivable that attempts to achieve the same with any F_ST_ outlier tests in data with this high neutral background level of differentiation would likely fail (e.g., Hoban et al. 2016; Liet al. 2019). Besides, as pointed out by Bierne *et al*. (2013), outlier tests are not expected to work well when genetic architecture of local adaptation has a polygenic basis, as is likely to be the case for most quantitative traits of ecological interest.

While the differences in head size and shape were very clear and consistent between marine and freshwater populations, the driftsel analysis revealed one anomaly: head traits in one of the pond populations (KRK) seemed to diverge in an orthogonal direction from marine and pond populations. While it might be tempting to ascribe such a deviation to random genetic drift from the hypothetical ancestor, it is also possible that there might be a biological explanation for this. Namely, in earlier analyses of behavioural differences between the same marine and freshwater populations, the KRK population showed a similar deviation in response to predation risk by piscine predators (Fraimout et al. 2021). What differentiates the KRK population from other studied pond populations is that it has a history of artificial introduction of brown trout (*Salmo trutta*) to this locality (Herczeg et al. 2009, 2010). Hence, while sticklebacks from the three other pond populations have evolved in absence of piscine predators since the ponds became isolated, the fish in KRK have been faced with recent brown trout predation. Hence, it is conceivable that this might have selected their behaviour and head morphology to converge back towards that of marine ancestors. Nonetheless, no piscine predators were observed in the field at the time of sampling and further studies should investigate the link between predation pressure and head shape morphology in *P. pungitius*.

While the evidence for adaptive differentiation in head size and shape was clear-cut, the results regarding the gill-raker morphology were not. We observed significant genetically based differentiation in gill-raker number and length between pond and marine fish, this differentiation did not exceed neutral expectations. This result does not exclude the possibility that natural selection could be behind the observed differentiation, only that statistical evidence for this is lacking. The consistent results for head gill-raker traits confirmed that the pond fish have evolved to become better suited for feeding on planktonic prey (longer snouts, upturned jaws, more and longer rakers) and marine fish better at feeding benthic prey with a blunt head, short snouts, fewer and shorter rakers. While this might at the first sight seem counterintuitive and go against expectations grounded on the work done in the related three-spined stickleback in which freshwater fish have benthic like heads and gill-rakers and marine fish limnetic like heads and gill-rakers (Rundle et al. 2003; Reimchen and Nosil, 2006; Raeymaekers et al. 2007), one has to keep in mind that there are subtle differences in ecologies of the three- and nine-spined stickleback species (Merila 2013). Specifically, the lack of piscine predators in isolated ponds opens up the pelagic niche for nine-spined sticklebacks (Karlsson & Byström 2005) and could favor development of more numerous and longer gill-rakers. Conversely, marine nine-spined sticklebacks (and those living in large lakes with predatory fish) living with piscine predators differ from three-spined sticklebacks in that they are not as gregarious as three-spined sticklebacks and tend to hide and feed in protection vegetation. Hence, the lack of marine three-spined stickleback-like gill-raker morphology in marine nine-spined sticklebacks makes sense if they rely less on planktonic diet and feed more benthically compared to syntopic three-spined sticklebacks.

What would be the selective advantage of the large head with strongly upward protruding jaw for pond sticklebacks? In absence of data on diet composition and/or trophic positioning data derived from stable isotopes from the study populations, we can only speculate to this effect. In the absence of native predatory fish, sticklebacks colonising freshwater habitats tend to evolve large heads with strongly upward protruding jaws (Walker & Bell 2000; Bell & Aguirre 2013; but see: Voje et al. 2013; Østbye et al. 2016). Such head morphology is thought be an adaptation to feed on limnetic diets and improved feeding performance on zooplankton (Lavin & McPhail 1986; Walker & Bell 2000). On the other hand, the head shapes of freshwater three-spined sticklebacks feeding primarily either on a limnetic or benthic diet (Wootton 1984; Gow et al. 2007) resemble closely those of pond and marine nine-spined sticklebacks, respectively. This could indicate that pond nine-spined sticklebacks released from predation pressure from piscine predators have shifted their diet towards feeding more zooplankton. Data from one of our study ponds (Rytilampi) indicates that although sticklebacks in this pond tend to feed mostly benthic prey, they do also consume pelagic zooplankton (Merilä & Eloranta 2017). Furthermore, data from Swedish lakes with predatory fish show that large nine-spined sticklebacks tend to be more planktivorous than their smaller conspecifics as they experience reduced predation risk in the pelagic open-water areas (Karlsson & Byström, 2005). The results of the analyses of gill-raker morphology align with these inferences: pond sticklebacks had long gill-rakers typical for fish adapted on feeding zooplankton, whereas marine ones had gill-rakers typical for benthivorous fish (Ingram et al. 2012). Hence, while the information gleaned from literature gill-rakers suggests that the observed divergence - clearly driven by divergent natural selection - is likely associated with differences in food acquisition, further studies looking into diets and feeding modes are needed to establish this firmly. Nevertheless, what is clear is that the observed divergence in head morphology cannot be explained by neutral process or phenotypic plasticity, and that the observed differences are not subtle, but apparent also for the naked eye (Fig.1b) and corresponding to substantial differences with large effect sizes.

The results further confirm that not only are freshwater stickleback populations adapted to their respective habitats, but also possess ample heritable variation in both relative head size and shape. Heritability estimates were fairly similar for marine and pond populations, suggesting that the latter have not suffered from massive loss of adaptive variation in spite of having lost significant amounts of neutral genetic variation due to population size bottlenecks and drift (Shikano et al. 2010; Kivikoski et al. 2023). Although these heritability estimates derive from full-sib analyses and hence potentially include maternal effect and dominance variance, earlier analyses of body size variation from two of these populations suggest that maternal effects dissipate quickly (Shimada et al. 2011) and there are no quantifiable dominance effects (Ab Ghani et al. 2012; Fraimout et al. 2021). Hence, we conclude that head size and shape are heritable, and hence able to respond to natural selection in all studied populations also in future.

Finally, although the mean head size and shape differed between marine and pond populations, one should note that there was also some degree of overlap among pond and marine populations, and heterogeneity in mean values of these variables as for instance reflected in variance in the degree that the different ponds were indicated to have diverged from the reconstructed ancestral form. This is not surprising given that freshwater habitats are heterogenous ecosystems in terms of their biotic and abiotic conditions and therefore, the direction and strength of natural selection on phenotypic traits are unlikely to be equal in all ponds. In fact, this kind of heterogeneity in mean trait values in replicate freshwater populations of sticklebacks has been seen also in earlier studies (e.g., Kaeuffer et al. 2012; Østbye et al. 2016).

In line with earlier results (Herczeg et al. 2010), we observed strong sexual dimorphism in head size and shape, males having larger differently shaped heads than females. While this might be indicative of niche partitioning among the sexes, it is also possible that sex differences are related to different sex roles. Nine-spined sticklebacks males use their jaws to build nests into which females lay their eggs, and males also defend their nests and hatchling offspring with their mouths (Wootton 1984). The degree of divergence among the two sexes was less compared to that among habitats - this may suggest that strength of habitat associated selection exceeds that of sex-specific selection.

### Conclusions

To sum up, the results provide evidence for adaptive divergence in head size and shape between pond and marine nine-spined sticklebacks, as well as genetically based divergence in gill-raker length and number. However, we failed to find evidence to indicate that the latter differentiation has been driven by natural selection. We further show the male and female sticklebacks display pronounced sexual dimorphism in head size and shape, and that further studies would be needed to understand the functional significance of this dimorphism. The result further suggests moderate to high heritability of relative head size and head shape suggesting that these traits have high potential to evolve in response to natural selection also in future.

## Ethical statement

All experiments were conducted under a permit from the Animal Experiment Board in Finland (permit reference ESAVI/4979/2018). Permission to collect fish from PYO, KRK and RYT ponds was obtained from Metsähallitus (license # MH794/2018). Authorisation to catch fish from the marine populations was provided with personal National Fishing Licenses.

## Data Availability

The raw data underlying this article including pictures, TPS files and genotype data will be made available in the Dryad Digital Repository. R scripts to replicate all analyses are publicly available at: https://github.com/afraimout/

## Author contributions

J.M. and A.F. conceived and designed the research. A.F. conducted field work and collected field samples. A.F. and Y.C. performed wet lab and data analyses. J.M. and A.F. led the writing effort, K.R. and Y.C. provided input and reviewed the manuscript.

## Acknowledgements

We thank Jacquelin DeFaveri, Niko BjoCrkell, Karlina Ozolina, Niina Nurmi, and Miinastiina Issakainen for their help in sampling and rearing sticklebacks. M. Issakainen also helped with DNA extractions. LEE Jae Hyun kindly drew the gill raker diagram. We thank Michael Bell for guidance on staining and measuring gill rakers. Our research was funded by the Academy of Finland (grant # 218343 to JM). The authors have no conflict of interest to declare.

## Notes

https://github.com/afraimout/

### Competing Interest Statement

The authors have declared no competing interest.

