## Supplemental figure 1 for "Divergent evolution of head morphology between marine and freshwater sticklebacks"

**Supplementary material for: Divergent evolution of head morphology between marine and freshwater sticklebacks**

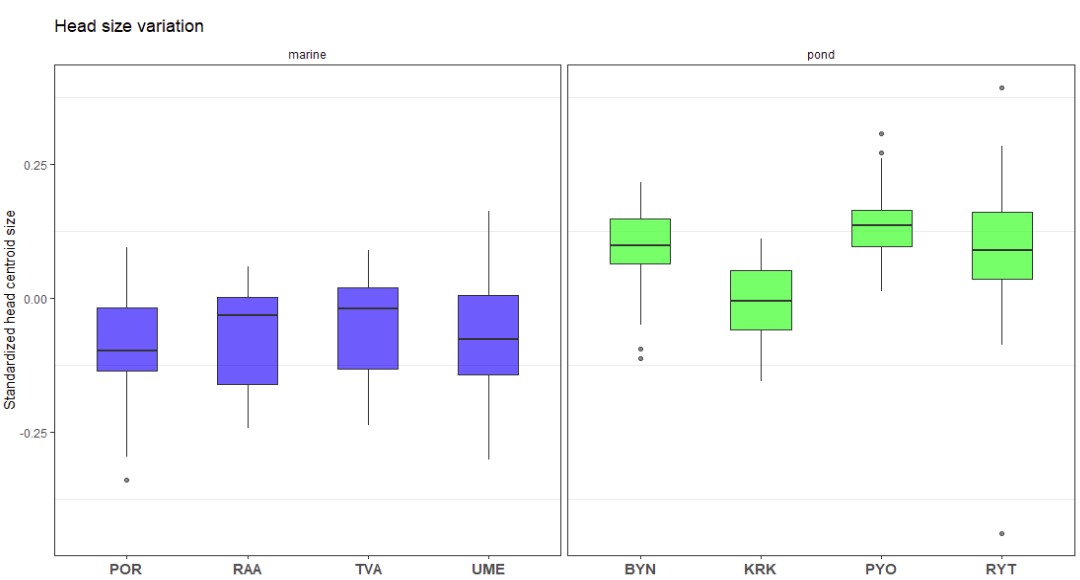

**Figure S1. Population differences in head centroid size.** Differences in head centroid size standardized for body length are shown for each habitat (marine: blue; pond: green) and populations.
